# Thumb-domain dynamics modulate the functional repertoire of DNA-Polymerase IV (DinB)

**DOI:** 10.1101/2022.10.16.512411

**Authors:** Damasus C. Okeke, Jens Lidman, Irena Matečko-Burmann, Björn M. Burmann

**Affiliations:** Department of Chemistry and Molecular Biology, University of Gothenburg, 405 30 Göteborg, Sweden; Wallenberg Centre for Molecular and Translational Medicine, University of Gothenburg, 405 30 Göteborg, Sweden; Department of Psychiatry and Neurochemistry, University of Gothenburg, 405 30 Göteborg, Sweden

**Author notes:** Correspondence should be addressed to BMB.

## Abstract

In order to cope with the risk of stress-induced mutagenesis, cells in all kingdoms of life employ Y-family DNA polymerases to resolve resulting DNA lesions and thus maintaining the integrity of the genome. In *Escherichia coli* (*E. coli*) the DNA polymerase IV, or DinB, plays this crucial role in coping with these type of mutations *via* the so-called translesion DNA synthesis. Despite the availability of several high-resolution crystal structures important aspects of the functional repertoire of DinB remain elusive. In this study, we use advanced solution NMR spectroscopy methods in combination with biophysical characterization to elucidate the crucial role of the Thumb domain within DinB’s functional cycle. We find that the inherent dynamics of this domain guide the recognition of double-stranded (ds) DNA buried within the interior of the DinB domain arrangement and trigger allosteric signals through the DinB protein. Subsequently, we characterized the RNA polymerase interaction with DinB, revealing an extended outside surface of DinB and thus not mutually excluding the DNA interaction. Altogether the obtained results lead to a refined model of the functional repertoire of DinB within the translesion DNA synthesis pathway.

## INTRODUCTION

Genomic DNA is constantly exposed to DNA-damaging toxins, radiations and genotoxic metabolites that cause DNA lesions or abnormal adducts within DNA strands ^1^. The Y-family DNA polymerases specialize in bypassing a wide range of DNA lesions. These polymerases function by accommodating aberrant DNA bases and perform nucleotide incorporation directly opposite the lesion, in a process known as translesion DNA synthesis (TLS) ^2^. Members of the Y-family polymerases are highly substrate specific ^3^, and possess high translesion synthesis accuracy, despite low fidelity on normal DNA synthesis, low catalytic efficiency, and low processivity. They also lack 3’ – 5’ proofreading exonuclease activity, and depending on the nature of the lesion may be error-prone or error-free ^4^. Evolutionary studies identified polymerases belonging to this family across all domains of life, e.g., DNA polymerase IV and polymerase V in *Escherichia coli*, Dpo4 and Dbh in yeast, and DNA polymerase η, ι, κ, as well as Rev1 in eukaryotes ^3,5,6^. All these polymerases share a conserved three-dimensional structure, a right-handed fold with two functional regions, the catalytic core, and the extended regulatory region (usually called either the little finger (LF) or polymerase-associated domain, PAD). The catalytic core region on the other side is composed of the three subdomains: Thumb, Palm, and Fingers ^3,7–9^.

The *E. coli* DNA polymerase IV is known to be induced as part of the SOS response to DNA damage-induced stress and plays a major role in adaptive mutagenesis ^10^. Kenyon and Walker named the genes identified to be inducible by MMC as *din* (for damage-inducible), and the bacterial translesion DNA polymerase IV, encoded as *dinB* is one of those genes ^11^. It has been demonstrated that DinB can accurately bypass mitomycin C (MMC)-induced N^2^-furfurly-deoxyguanosine (N^2^-fdG) as well as ultraviolet (UV) radiation-induced thymine-thymine cyclobutane pyrimidine dimer (CPD) kind of lesions with high catalytic efficiency ^12,13^. DinB shares the catalytic core – regulatory region structural architecture common to Y-family polymerases, which could be established by several crystallographic studies ^14^. DinB is able to bind to the DNA substrate *via* the Thumb domain with the PAD providing an additional interface ^15^. The Palm domain coordinates the two Mg^2+^ ions through the carboxylate group of Asp8 and Asp 102, and the Fingers domain interacts with the template DNA base and the incoming deoxynucleotide (dNTPs) substrates ^16^. The extended channel between the Fingers and the PAD provides an adequate space for the accommodation of bulky lesions, thereby preventing any steric hinderance as well as enabling their accurate and error-free bypass ^14^. The PAD has been demonstrated to mediate DinB recruitment to the site of stalled replicative DNA polymerase and towards transcribing RNA polymerase (RNAP) ^17,18^, an interaction proposed to be modulated by β-clamp processivity factor ^19,20^ and the NusA transcription elongation factor ^21^, respectively. Besides the β-clamp factor, the physiological role of DinB has been shown to be mediated by UmuD, UmuD’, RecA, and the molecular chaperone GroEL ^22–24^.

Crystallographic studies of *E. coli* DinB in complex with damaged double-stranded DNA and incoming nucleotide ^14^ reveal that the Thumb domain adopts a helix-hairpin-helix (HhH) motif consisting of three short helices separated by flexible loops. The amino-terminal end of the Thumb domain is linked to the Palm domain whilst the carboxy-terminal end is connected to a flexible extended linker linking the catalytic core to the PAD. The Thumb is the smallest domain within the catalytic core and located at the edge of the core, enabling conformational re-orientation of the domain to form a large cavity in the presence of bulky DNA substrate ^14^. Experimental evidence suggests that the catalytic core and the PAD freely move in relation to each other in the absence of DNA substrate and undergo large conformational change upon DNA binding ^25,26^. Despite the vital role of this domain in the accommodation of abnormal DNA bases in the DinB active site, the structural dynamics and its conformational adaptions for DinB and its related proteins remain elusive. Therefore, we chose to study DinB in solution to assess its dynamical properties *en detail.* Using advanced high-resolution solution-state NMR we provide intially sequence specific resonance assignment of *E. coli* DinB forming the foundation of in-depth studies of its inherent dynamics over a broad range of timescales as well as its structural adaptations upon DNA as well as RNAP binding. Employing Biolayer Interferometry (BLI) to study the different interactions permitted us to elucidate the key role of the Thumb region in facilitating these as well as confirming the auxiliary role of the PAD. Together our dynamic and interaction data enabled us to refine the functional picture of DinB, which likely has implications for the vast majority of its related proteins spanning all kingdoms of life.

## METHODS

### Cloning

The used DinB constructs were subcloned from a pET28b(+)-DinB construct (purchased from Genscript), yielding DinB^1-351^ with an amino-terminal His_6_-tag. The individual domain constructs (DinB-NTD (DinB^1–165^), DinB-Palm (DinB^1–10, 74–165^), DinB-Fingers (DinB^11–73^), DinB-Thumb (DinB^166–241^), and DinB-PAD (DinB^228–351^)) were cloned into a pET28_SUMO vector using a pET28_SUMO_Hsc70 plasmid (a kind gift of B. Bukau (Heidelberg)) as a template, yielding amino-terminally SUMO-tagged DinB constructs using standard methods. DinBΔPAD (DinB^1–230^) was cloned by introducing a stop codon at amino acid 231 using standard single-point mutagenesis. Plasmids and primers used in this study can be found in **Supplementary Tables S1**.

### Expression and Purification of Proteins

Each of the DinB constructs were chemically transformed into *E. coli* BL21 (λDE3) cells and subsequently grown in 1L LB (Luria Bertani) medium supplemented with 50 *μ*g/ml kanamycin at 37°C. Upon reaching an OD_600_ of 0.6 – 0.8 the cells were induced with 1 mM IPTG (isopropyl β-D-1-thiogalactopyranoside; Thermo Scientific) and expression was continued overnight (12–16 h) at 25°C. The cells were harvested by centrifugation at 5,000*g for 20 minutes at 4 °C, resuspended for lysis in 40 ml buffer A (50 mM HEPES, 500 mM NaCl, 5 mM Imidazole, 1 mM DTT, pH 7.5; ~1:4 ratio cell pellet weight), supplemented with 5 mM MgCl_2_ (final concentration), cOmplete EDTA-free protease inhibitors cocktail (Roche Diagnostics GmbH), and HL-SAN DNase I (ArticZymes), and incubated for 30 minutes at 4°C. Thereafter, the cells were lysed by three passes through an Emulsiflex C3 (Avestin) homogenizer at 4 °C. Cell debris was removed *via* centrifugation at 19,000*g for 50 min at 4°C. All different DinB variants were purified using a 5 ml Ni^2+^-Histrap HP column (GE Healthcare) column equilibrated with buffer A and subsequently eluted with buffer B (buffer A supplemented with 1 M imidazole). Eluted fractions of DinB and DinBΔPAD were dialyzed overnight against Buffer C (25 mM HEPES, 200 mM NaCl, 50 mM arginine, 50 mM glutamate, 2 mM DTT, 1 mM EDTA, 0.5 mM CHAPSO, pH 7.5), while His_6_-SUMO-variants were dialyzed against buffer D (25 mM HEPES, 200 mM NaCl, 2 mM DTT, pH 7.5), respectively. The His_6_-SUMO tag was subsequently cleaved with human SenP1 (Addgene #16356) ^27^ and separated on Ni^2+^-Histrap HP column. All proteins were further purified on a 5 ml HiTrap Heparin HP column (GE Healthcare), equilibrated in their respective dialysis buffer and eluted by an NaCl gradient containing 1 – 1.5 M NaCl. Eluted fractions from the Heparin step were pooled and concentrated using appropriate Vivaspin centrifugal concentrators (MWCO FF; Sartorius). DinB and DinBΔPAD, respectively, were additionally purified on a size exclusion chromatography column (Superdex 200 Increase column (GE Healthcare)) equilibrated with buffer E (50 mM KPi, 1 M KCl, 50 mM arginine, 50 mM glutamate, 10 mM MgCl_2_, 2 mM DTT, 1 mM EDTA, 0.5 mM CHAPSO, pH 7.0). All the purified proteins were concentrated to a range of 0.3 – 1 mM and stored −80°C until further usage.

### Isotope Labeling

Isotope labeling of proteins was achieved by expressing protein in 2x M9 minimal media ^28^ with the relevant isotopes supplemented to the media (H_2_O supplemented with (^15^NH_4_)Cl for [*U*-^15^N]-labeled protein or with both (^15^NH_4_)Cl and *D*-(^13^C)-glucose for uniform double labeling [*U*-^13^C,^15^N], or D_2_O supplemented with (^15^NH_4_)Cl yielding [*U*-^2^H,^15^N]-labeled proteins or *D*-(^2^H, ^13^C)-glucose for uniform double labeling [*U*-^2^H,^13^C,^15^N]). For specific methyl group labeling, Met, Ala, Leu, Val, and Ile (MALVI)-labeled DinB sample was produced in Bioexpress rich (2 %) D_2_O-based 2xM9 media supplemented with (^15^NH_4_)Cl, *D*-(^2^H,^12^C)-glucose as well as 50 mg/l 2-Ketobutyric acid-4-^13^C,3,3-d_2_ sodium salt hydrate (Isoleucine), 4 vials/l DLAM-LV^*proS*^-kit (2-(^13^C)-methyl-4-(D_3_)-acetolactate (valine/leucine *proS* methyl group only)), 50 mg/l [*U*-^2^H, ^13^CH_3_] methionine, and 50 mg/l 2-[^2^H], 3-[^13^C] L-alanine (alanine) precursors added 1 h prior to induction ^29–31^. Bioexpress, methionine, and alanine were purchased from Cambridge Isotopes Laboratories and the DLAM-LV^*proS*^ precursors from NMR-Bio. All other isotopes were purchased from Merck.

### DNA oligonucleotide preparation

Two self-complementary single-stranded DNA (ssDNA) oligonucleotides (18 mer: 5’-TCTAGGGTCCTAGGACCC – 3’ and 5’ – GGGTCCTAGGACCCTAGA – 3’) were purchased from Eurofins in lyophilized form. The oligonucleotides were resuspended in annealing buffer (10 mM Tris-HCl, 50 mM NaCl, 1 mM EDTA, pH 7.8) to a concentration of 340 *μ*M. To obtain double-stranded DNA (dsDNA), equimolar concentrations of the ssDNA were mixed together and heated for 5 minutes in a 95 °C heating block. Thereafter, the heating block was turned off to allow the ssDNA oligonucleotides anneal together while gradually cooling down to room temperature. The sample was exchanged into appropriate NMR buffer for protein-dsDNA NMR titration using Zeba™ Spin Desalting Columns (7K MWCO, Thermo Fisher Scientific).

### RNAP core enzyme expression and purification

RNA polymerase core was expressed from plasmid pIA900 (Addgene #104401) ^32^ and purified according to an established protocol ^32^ with the exception that the Mono Q^®^ ion-exchange purification step was exchanged with a size exclusion chromatography step using a Superose6 10/300 GL column (GE Healthcare) pre-equilibrated with PBS buffer ^33^ supplemented with 2 mM DTT. A sample of purified RNAP core was ran on SDS-PAGE to confirm the presence of all subunits. The concentrated protein was stored in −80°C until further usage.

### NMR Spectroscopy

NMR measurements were performed on Bruker Avance III HD 700 MHz, 800 MHz or 900 MHz spectrometers, running Topspin 3.5/3.6 and each equipped with a cryogenically cooled triple-resonance probe. All experiments were performed at 298 K in NMR buffer as listed in Table S3.

For the sequence-specific backbone resonance assignment of DinB and DinBΔPAD constructs, 2D [^15^N, ^1^H]-TROSY-HSQC ^34^ as well as the following TROSY-type 3D experiments: 3D trHNCA, 3D trHNCACB, and 3D trHNCO ^35^ experiments were recorded. Whereas for the different domain sub-constructs, standard through-bond 3D HNCA, 3D HNCACB, 3D HNCO, 3D HN(CA)CO, 3D CBCA(CO)NH ^36^ experiments were used. Aliphatic side-chain resonance assignment for the different sub-constructs were performed based on 2D [^13^C, ^1^H]-HSQC spectra with/without constant time (CT) version, as well as 3D (H)CC(CO)NH, H(CC)(CO)NH, and HCCH-TOCSY-experiments ^36^. In addition, the following NOESY-type experiments with the indicated mixing times were performed: 3D ^13^C_methyl_–^13^C_methyl_–^1^H_methyl_ SOFAST NOESY with 50 ms and 300 ms mixing time ^37^.

For the quantitative analysis of signal intensities, the amplitudes were corrected by differences in the ^1^H-90° pulse-length, the number of scans, and the dilution factor ^38^. NMR data were processed with a combination of Topspin 4.0.7 (Bruker Biospin), NMRPipe ^39^ and mddNMR2.6 ^40^ as well as analyzed with CARA ^41^.

Secondary chemical shifts were calculated relative to the random coil values using the prediction software POTENCI ^42^. Further, a weighting function with weights *1–2–1* for residues (*i-1*)–*i*–(*i+1*) was applied to the raw data ^43,44^.

For titration experiments, 2D [^15^N, ^1^H]-TROSY-HSQC experiments were acquired with 16 scans and 2048 × 256 complex points in the direct and indirect dimensions, respectively. The chemical shift changes of the amide moiety were calculated as follows:

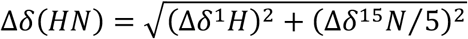

### Methyl group assignments

For the methyl group assignment we used a [*U*-^2^H, Ile-δ_1_-^13^CH_3_, Leu-δ_2_, Val-γ_2_-*proS*-^13^CH_3_, Ala-β-^13^CH_3_, Met-ε-^13^CH_3_]*–*DinB sample, termed MALVI^*proS*^-DinB, with only the *proS* methyl group of the valine and leucine group labeled ^29^. We took advantage of the available methyl group assignments obtained already by the resonance specific assignment of its subdomains DinB-PAD, DinB-Thumb, and DinBΔPAD obtained in this work. To resolve ambiguities within the MALVI^*proS*^-DinB sample we used a 3D ^13^C_Methyl_–^13^C_Methyl_–^1^H_Methyl_ SOFAST NOESY experiment with mixing times of 50 ms and 300 ms. This approach yielded the following degree of assignment (~93%): Alaβ (31/31), Ileδ_1_ (22/22), Leuδ_2_ (33/38), Metε (9/10), and Valγ2 (21/24).

### NMR backbone dynamics

For the analysis of the dynamic properties of DinB, the following relaxation experiments were measured: ^15^N{^1^H}-NOE, *T*_1_(^15^N), and *T*_1ρ_(^15^N) ^45^. Non-linear least square fits of relaxation data were done with Pint ^46^. *R*_2(R1ρ)_(^15^N) values were derived from *T*_1ρ_ using equation 2:

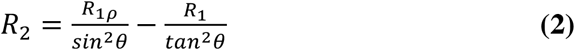

with 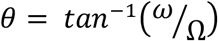, where ω is the spin-lock field strength (2 kHz) and Ω is the offset from the ^15^N carrier frequency ^44^.

Error bars for *R*_1_(^15^N) and *R*_1ρ_ (^15^N) were calculated by a Monte Carlo simulation embedded within Pint ^46^, and for *R*_2(R1ρ)_ (^15^N) by error propagation. Error bars for the ^15^N{^1^H}-NOE were calculated from the spectral noise. Analysis of the obtained relaxation rates was performed using an anisotropic diffusion tensor using TENSOR2 ^47^ on the NMRbox web server ^48^.

### NMR side chain dynamics

Experiments were performed on a [*U*-^2^H, Ile-δ_1_-^13^CH_3_, Leu-δ_2_, Val-γ_2_-*proS*-^13^CH_3_, Ala-β-^13^CH_3_, Met-ε-^13^CH_3_]*–*DinB sample, with only the *proS* methyl group of the valine and leucine group isotopically labeled ^29^, at a temperature of 25°C in 99.9% D_2_O based NMR buffer. Side chain methyl order parameters (S^2^_axis_) were determined by cross-correlated relaxation experiments ^49,50^. Single- (SQ) and triple-quantum (TQ) ^1^H–^13^C experiments were collected at a series of delay times. Ratios of the peak intensities were fitted for four values ranging between 2 and 16 ms using the following equation where *T* is the relaxation delay time and δ a factor to account for coupling due to relaxation with external protons:

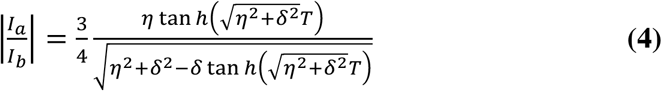

S^2^_axis_ values were determined using equation 5:

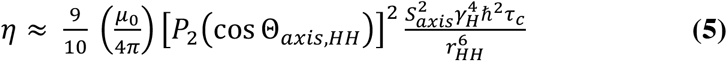

Where *μ*_0_ is the vacuum permittivity constant, γ_H_ the gyromagnetic ratio of the proton spin, *r*_HH_ is the distance between pairs of methyl protons (1.813 Å), S^2^_axis_ is the generalized order parameter describing the amplitude of motion of the methyl 3-fold axis, Θ_axis,HH_ is the angle between the methyl symmetry axis and a vector between a pair of methyl protons (90°), and P_2_(x) = ½ (3x^2^-1). Data was analyzed by in-house written scripts and finally, the product of the methyl order parameter and the overall correlation time constant, S^2^_axis_•τ_C_, was determined.

Multiple quantum (MQ) methyl relaxation dispersion experiments ^51^ were recorded as a series of 2D data sets using constant time relaxation periods (*T*) of 40 ms (800 MHz) and CPMG (Carr-Purcell-Meiboom-Gill) frequencies of 25 and 750 Hz. *R*_2,eff_, the effective transverse relaxation rate was calculated according to the following equation:

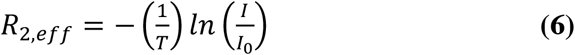

where *I* (or *I*_0_) are the intensities with and without the presence of a constant-time relaxation interval of duration *T*, during which a variable number of ^13^C 180° pulses are applied leading to ν_CPMG_ = 1/(2δ), where d is the time between successive pulses. Data were analyzed by Pint ^46^ to extract signal intensities and determineΔ*R*_2,eff_, the difference between the rates at 25 Hz and 750 Hz.

### Bio-Layer Interferometry (BLI)

BLI experiments were performed on an Octet RED96 system (Fortébio) at 303 K as outlined *en detail* before ^33,52^. Briefly, ligands (DinB, DinBΔPAD, and DinB-PAD) were biotinylated using the biotinylation kit EZ-Link NHS-PEG4-Biotin (Thermo Fisher Scientific). The biotin label was freshly dissolved in H_2_O, directly added to the protein solution in a final molar ratio of 1:1 in PBS (pH 7.4) buffer supplemented with 1 mM TCEP followed by gentle mixing at room temperature for 45 min. The used reaction conditions favored the preferential labeling of the amino-terminal α-amino group of proteins ^53^. Unreacted biotin was removed with Zeba™ Spin Desalting Columns (7 MWCO, Thermo Fisher Scientific). The kinetic assay between DinB variants and RNAP core was carried out by immobilizing biotin-labeled proteins (0.5 μM Biotin-DinB, 0.5 μM Biotin-DinBΔPAD or 1 μM Biotin-DinB-PAD) onto hydrated streptavidin (SA) biosensors. Subsequently, the ligand-bound biosensors were blocked with EZ-Link Biocytin (Thermo Fisher Scientific). Then, RNAP core (analyte) was serially diluted in 2-folds and applied to the ligand-bound biosensors in 50 mM KPi, pH 6.5, 150 mM NaCl, 25 mM MgCl_2_, 1 mM TCEP, 0.5 % BSA (for DinB – RNAP core assay), 50 mM KPi, pH 6.5, 50 mM NaCl, 25 mM MgCl_2_, 1 mM TCEP, 0.25 % BSA (for DinBΔPAD – RNAP core assay), and 50 mM KPi, pH 6.5, 300 mM NaCl, 1 mM TCEP (for DinB-PAD – RNAP core assay).

For the kinetic assay between DinB variants and dsDNA, biotin-labeled ssDNA oligonucleotide and its complementary strand were obtained directly from Eurofins and annealed as outlined in the DNA oligonucleotide preparation section above. Biotin-labeled dsDNA immobilization and ligand-bound biosensor blocking procedures were as stated above. Each of the DinB variants were serially diluted 2-folds and applied to the dsDNA-bound biosensors in 50 mM KPi, pH 6.5, 50 mM NaCl, 10 mM MgSO_4_, 1 mM TCEP. Double referencing (Sensor reference and Sample reference according to the manufacturer’s instructions) was used in all the assays to eliminate any background signal and/ or signal due to non-specific binding of analyte to the biosensor. The experiments were set up and the acquired data was subsequently analyzed using the Data acquisition 10.0 and the Data analysis HT 10.0 (Fortébio) software, respectively.

### Data availability

All data needed to evaluate the conclusions in the paper are present in the paper and/or Supplementary Materials. The sequence-specific NMR resonance assignments of DinB and the diverse subconstructs have been deposited in the BioMagResBank (BMRB) with accession codes XXX, YYY, ZZZ, respectively. The NMR data used for chemical shift perturbations, relaxation analysis have been tabulated and were deposited on Mendeley Data: UUU. Additional data related to this paper might be requested from the corresponding author.

## Results

### DinB structure in solution

As the structural information about DinB is thus far limited to static X-ray structures ^14^ and insight into its inherent protein dynamics remains largely elusive, we employed advanced high resolution NMR spectroscopy in solution to address the dynamic properties of DinB. To this end, we initially expressed the multidomain protein (**Figure 1A, B**) as uniformly labeled [*U*-^2^H,^15^N]-DinB and measured [^15^N,^1^H]-NMR spectra. These spectra were of excellent quality despite the size of ~40 kDa of DinB, indicating clearly a well folded protein with the expected number of resonances, comparable to the only previous NMR study of a DinB-homologue from *Sulfolobus acidocaldarius* ^54^. Despite these promising first results, we realized once using standard transverse relaxation-optimized spectroscopy (TROSY)-type three dimensional (3D) experiments, that complete resonance specific assignment, due to ambiguity as well as missing signals in some of the 3D experiments, would not be possible. To circumvent this issue, we resorted to a divide-and-conquer approach exploiting the modular architecture of DinB. Therefore, we cloned and purified the individual DinB domains. Despite several efforts and rounds of testing different expression constructs, we could not successfully express and purify the amino-terminal constructs of DinB, namely DinB-NTD, DinB-Palm and DinB-Fingers (**Figure 1A**). Thus, we put our focus on the three remaining constructs DinBΔPAD, DinB-Thumb and DinB-PAD (**Figure 1A**), which all yielded well dispersed and high quality [^15^N,^1^H]-NMR spectra (**Figure 1C**). Using standard triple resonance 3D experiments with [*U*-^13^C,^15^N]-PAD and [*U*-^13^C,^15^N]-Thumb yielded 100% and 84% complete sequence-specific backbone resonance assignment for the two constructs, respectively. Due to its larger size, we used a [*U*-^2^H,^13^C,^15^N]-DinBΔPAD sample together with TROSY-type 3D experiments yielding 95% complete sequence-specific backbone assignment. With the data of the different constructs, we were able to transfer and validate the assignments of full-length DinB reaching a final assignment completeness of ~95% for the backbone resonances.

**Figure 1.**
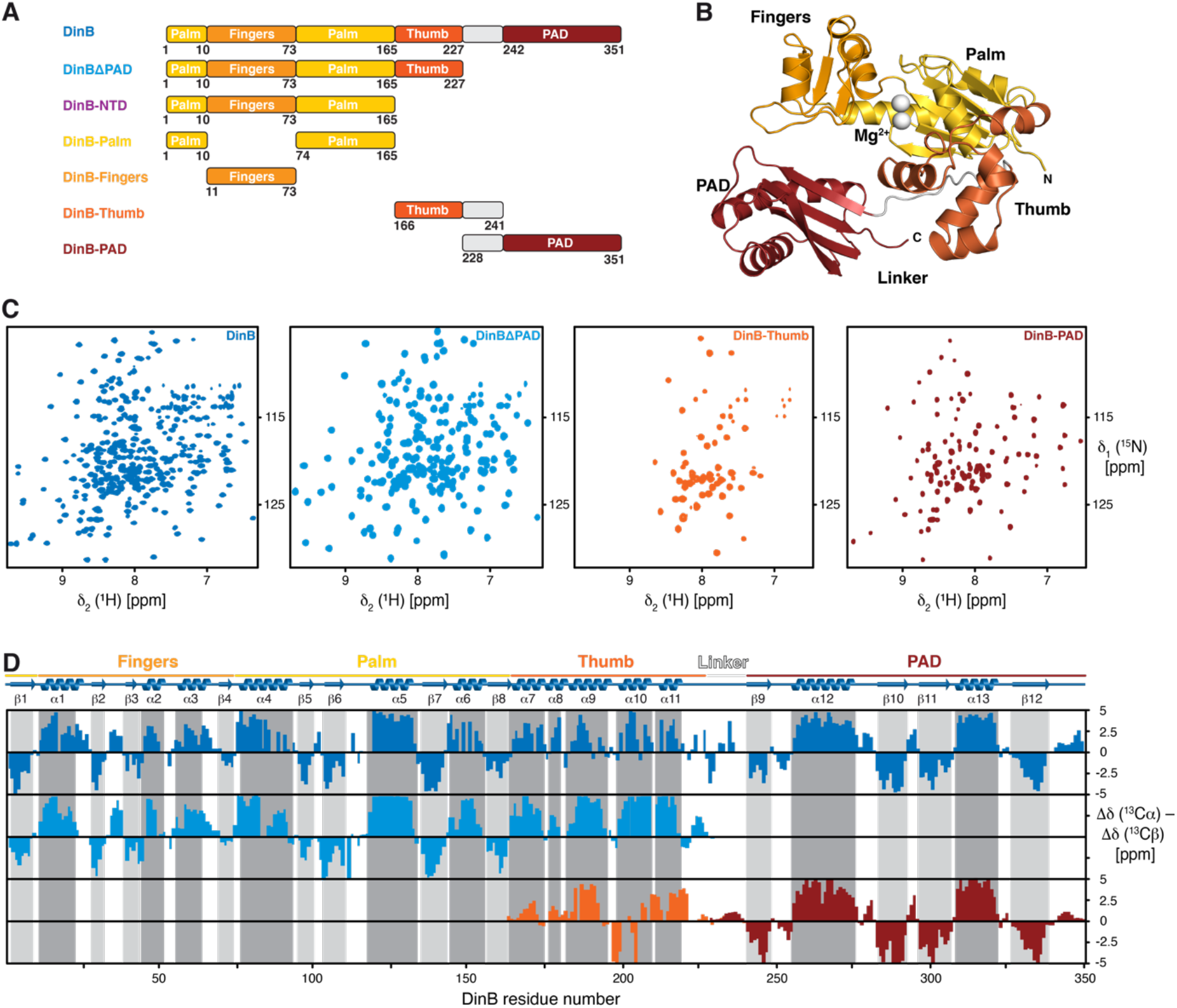
**A**) Scheme of the used constructs indicating the domain-structure of the DinB. **B**) Crystal structure of the *E. coli* DNA polymerase IV (PDB-ID: 4Q45) with the Fingers (orange), Palm (yellow), Thumb (orange-red) and PAD (dark-red) domains as well as magnesium ions (white) indicated. **C**) 2D [^15^N, ^1^H]-NMR spectra of the [*U*-^2^H,^15^N]-DinB, [*U*-^2^H,^15^N]-DinBΔPAD, [*U*-^15^N]-DinB-Thumb, and [*U*-^15^N]-DinB-PAD measured in NMR-buffer at 298K. **D**) Secondary structure elements of the DinB-domains in solution (color) as derived from the secondary backbone ^13^C chemical shifts. The secondary structure elements of DinB within the X-ray structure (PDB-ID:4Q45) are indicated in grey (β-strands) and dark-grey (α-helices).

Using the obtained resonance assignments for the different DinB constructs, we compared the combined ^13^Cα and ^13^Cβ chemical shifts to determine the secondary structure elements in solution and compared these to the crystal structure (**Figure 1D**). Whereas for DinB, DinBΔPAD and the isolated PAD the agreement was very good, and the individual elements might be shifted by a single residue only, the difference for the Thumb was more obvious. In addition, the extend of the secondary chemical shifts was less pronounced for this domain, indicating that individual α-helices such as α7 might only be populated for about 30% of the time pointing to some structural instability of this domain. Comparing this data to the full-length crystal structure (**Figure 1B**) clearly shows that the Thumb packs against a surface on the Palm region, which might stabilize this domain within the full-length protein. In accordance with this hypothesis, we also observed significant chemical shift changes of the amide moieties when comparing the isolated Thumb and the Thumb within the DinBΔPAD construct. Otherwise, the rather weak overall signal intensity of this domain within the larger DinB constructs points also to underlying dynamics on the fast to intermediate NMR timescale with exchange rates ~100 – 1000 s^−1^ in this region.

### DinB backbone dynamics

To address the conformational dynamics of DinB *en detail*, we evaluated in a next step the backbone dynamics of DinB over a broad range of timescales by NMR relaxation measurements ^45,55^. By measuring the steady-state heteronuclear ^15^N{^1^H}-NOE (hetNOE) and the ^15^N longitudinal (*R*_1_) relaxation rates we probed the pico- to nanosecond motions of the H–N bonds. In these type of experiments, high hetNOE values and small *R*_1_ rates indicate rigid and stably folded regions whereas the inverse, low hetNOE values as well as high *R*_1_ rates, points to flexible and unfolded segments. Consistent with our structural characterization, the hetNOE data indicated that the folded parts of the Fingers, Palm and PAD are stably folded as evident by high hetNOE values, whereas linker and loop regions are more flexible as indicated by low hetNOE values (**Figure 2A, B** and **Supplementary Figure 1**). In slight contrast was the observed behavior of the Thumb domain, which showed on average lower hetNOE values of 0.66 for the residues comprising helices, compared to 0.73 for the full-length protein, which in general indicates a stable fold. Overall, the obtained values for the DinB overall are well within the theoretical maximum expected at 18.8 T (800 MHz ^1^H frequency) of 0.86, indicating the presence of global fast motions within the whole domain in solution. These motions are most pronounced within the Thumb domain as this region shows the lowest average values, which might also stem from inherent dynamics as this domain also showed quite distinct chemical shift changes between separated form and within the full-length protein as discussed above.

**Figure 2.**
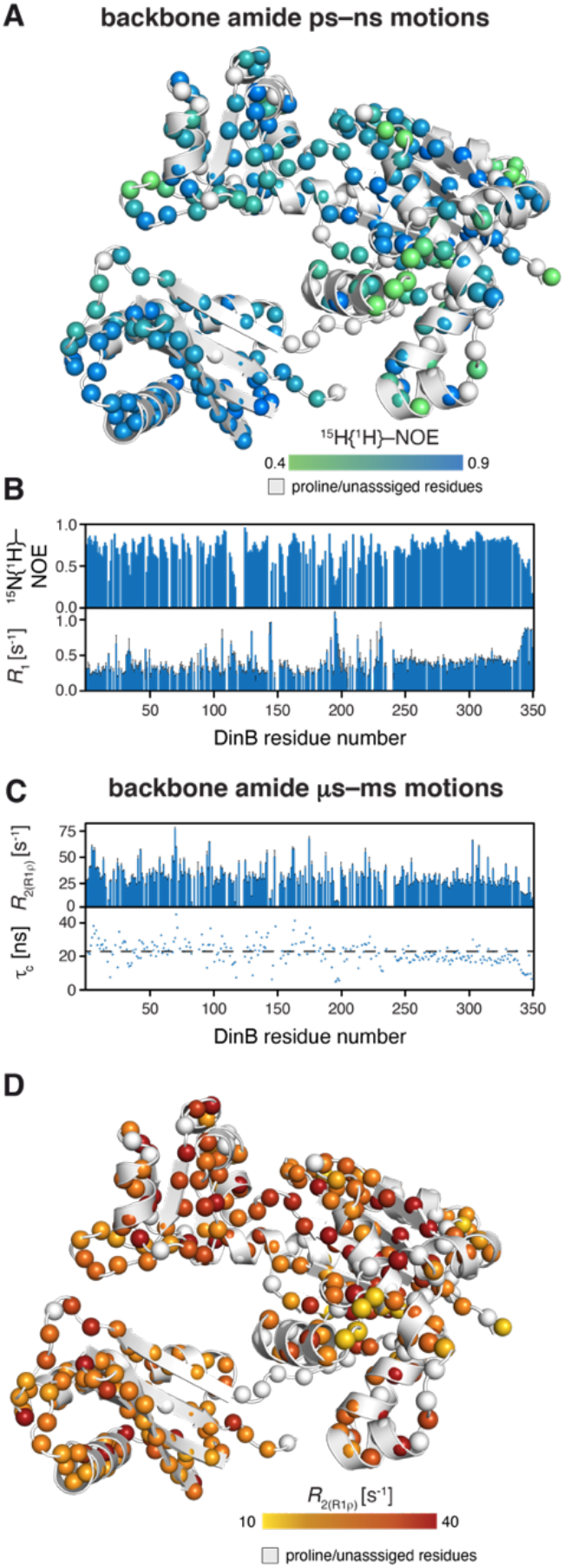
**A**) Dynamics on the ps–ns timescale plotted on the DinB structure (PDB-ID: 4Q45). The amide moieties are shown as spheres and the hetNOE values are indicated by the green to blue gradient. **B**) The hetNOE (top) as well as the longitudinal relaxation rate *R*_1_ are plotted against the DinB residue number. **C**) The *R*_2(R1ρ)_ (top) as well as the rotational correlation time τ_c_, obtained from the analysis of the *R*_1_ and *R_2_*_(R1ρ)_ relaxation rates, are plotted against the DinB residue number. The broken line indicates the average value of 21.5 ns. **D**) Dynamics on the *μ*s–ms timescale plotted on the DinB structure (PDB-ID: 4Q45). The amide moieties are shown as spheres and the *R*_2(R1ρ)_ values are indicated by the yellow to red gradient. Relaxation data was measured at 18.8 T.

The obtained average *R*_1_ rates for the folded segments of DinB could be determined to be 0.36 s^−1^, in line with the magnitude of the obtained hetNOE values. We observe only a marginal increase for the Thumb domain to 0.38 s^−1^ and a more notable one for the carboxy-terminal residues 342–351 with 0.76 s^−1^, indicating that this segment is intrinsically unstructured.

In a next step, we analyzed the contributions of motions in the micro- to millisecond regime. To assess these slow timescale motions, we analyzed the ^15^N transverse relaxation rates. We measured the *R*_2_-rates derived from the *R*_1ρ_ rates (*R*_2(R1ρ)_), which report on the motions on the lower micro-second timescale, because under the used spin-lock radio frequency (RF) field of 2,000 Hz all exchange contributions (*R*_ex_) much slower than 80 *μ*s would be leveled out. In line with the previous analysis of the fast timescale motions, we observed a largely planar profile for the folded segments with an average value of 29 s^−1^ compared to almost identical 28 s^−1^ for the Thumb domain (**Figure 2C, D** and **Supplementary Figure 1**).

Based on the *R*_1_ rates as well as the *R*_2(R1ρ)_ rates we next determined the rotational correlation time τ_C_ of the DinB with ~22 ns for the structured region (**Figure 2C**), which is in line for a protein of 40 kDa. We could observe that the values for the DinB-PAD were generally lower as for the rest of the well-structured parts of DinB, which was due to slightly altered *R*_1_ and *R*_2(R1ρ)_ rates. To assess if this modulation of the relaxation rates can be attributed to different global dynamics of the DinB-PAD, we analyzed next the *R*_1_/*R*_2(R1ρ)_ quotient as the distributions provide direct insight into if the analyzed domains tumble as (partly) independent units ^56,57^. Our data clearly showed a bimodal distribution (**Supplementary Figure 1**), which is a clear indication that in the substrate-free form the DinB-PAD is partially decoupled from the rest of the protein, in line with the supposed role of this domain.

In summary, a picture emerges where the highly flexible linker region connects a stably folded PAD to the rest of the DinB protein, whereas the Thumb domain exhibits a large scale of fast timescale motions on the pico- to nanosecond timescale, which could be either attributed to inherent flexibility or the priming for interactions with the DNA substrate.

### Side-chain dynamics elucidate the central role of the Thumb

To obtain a more detailed picture of the underlying dynamics within DinB and especially within its Thumb domain, we exploited the increased sensitivity of methyl groups to get access to the side-chain dynamics of the DNA polymerase IV. We chose specific labelling of isoleucine, alanine, and methionine, as well as stereo-specific labeling of valine and leucine methyl groups as these are well dispersed among the whole DinB, proving specific probes (**Figure 3A**). The exceptional quality of the obtained 2D [^13^C,^1^H]-NMR spectrum enabled us to assign ~93% of all methyl groups of the MALVI^*proS*^-DinB sample in a sequence specific manner (**Figure 3A**; details are provided in the Methods section).

**Figure 3.**
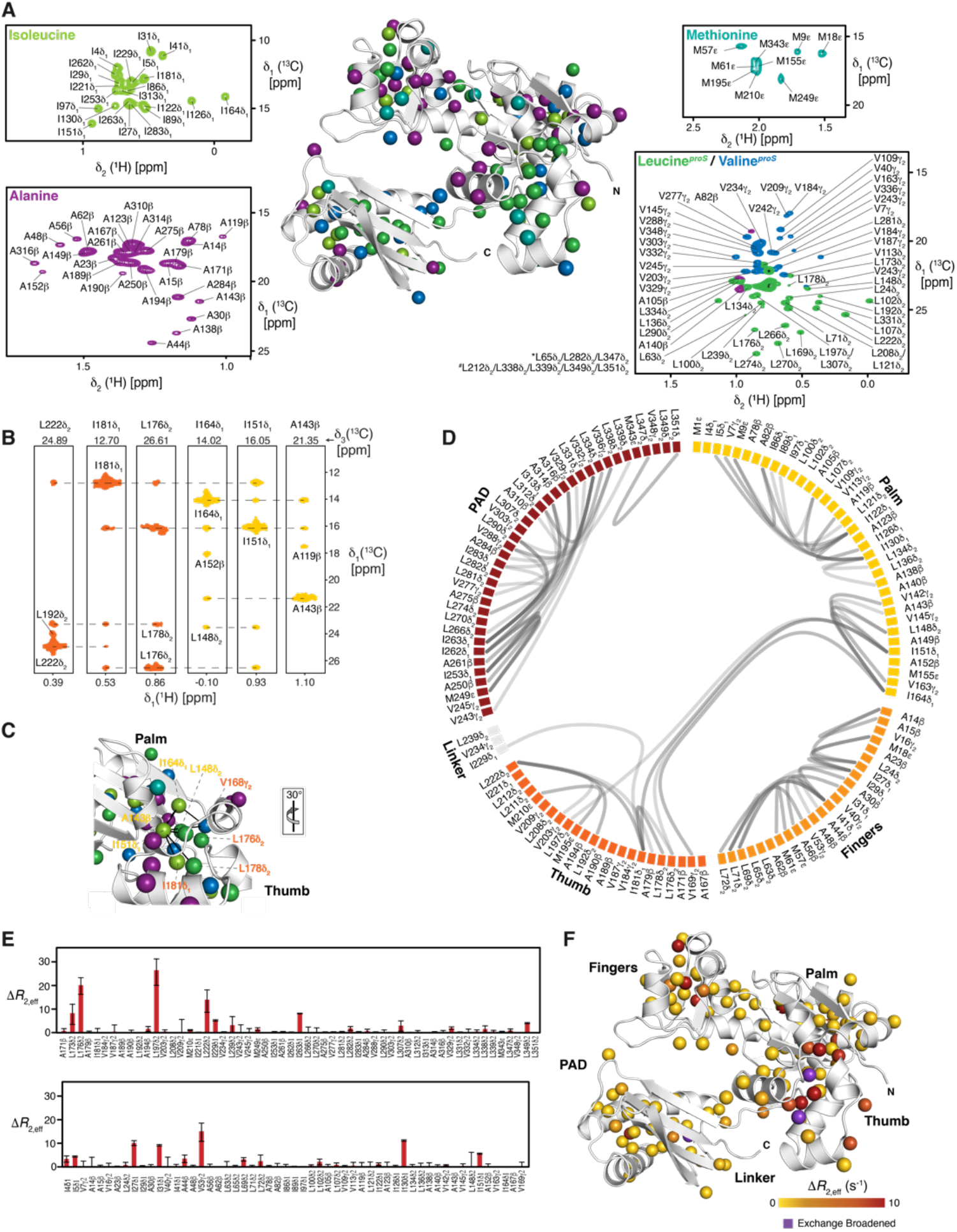
**A**) Distribution and assignment of isoleucine, alanine, methionine, leucine and valine methyl groups using an [*U*-^2^H, Ile-δ_1_-^13^CH_3_, Leu-δ_2_-^13^CH_3_, Val-γ_2_-^13^CH_3_, Ala-^13^CH_3_, Met-^13^CH_3_] stereospecific labelled DinB measured in NMR buffer at 298 K. **\*** and ^***#***^ denote leucine residues in the overlapping central region. **B)** Representative NOE strips from a 3D ^13^C_methyl_-^13^C_methyl_-^1^H_methyl_ SOFAST NOESY focusing on the interdomain stabilization between Palm (yellow) and Thumb (orange). **C)** NOE network, indicated by the black lines, stabilizing the Thumb on the Palm surface plotted on the DinB structure (PDB-ID: 4Q45). The respective orientation relative to panel **A** is indicated. **D)** Flareplot visualization of the complete methyl-methyl NOE network detected for the MALVI^*proS*^-DinB illustrating the connectivity between the individual domains **E, F**) Δ*R*_2eff_ values for the methyl-groups obtained from the difference of *R*_2,eff_ at the lowest and highest CPMG frequency υ_CPMG_ (**E**). Structural view of the amplitude of the CPMG relaxation dispersion profiles Δ*R*_2eff_ at 18.8 T (**F**). Exchange broadened residues are indicated

We initially characterized the methyl-methyl NOEs of this MALVI^*proS*^-DinB (**Figure 3B–D**). These NOE distances revealed a large network of contacts mainly within the individual domains. The only clear indication of domain–domain interaction *via* methyl-groups could be identified between the Palm and the Thumb, which is mainly facilitated by three central isoleucine residues, namely I151, I164, and I181 (**Figure 3B, C**).

Assessing the dynamics of the methyl-groups, we next determined the product of the side-chain order parameters and the correlation time of the overall molecular tumbling (S^2^_axis_•τ_C_), which reports on the extend of the amplitude of motions on the fast NMR timescale (**Supplementary Figure 2**) ^49,50^. The obtained values showed a maximum of ~23 ns, which is in good agreement with the determined τ_c_ of DinB based on the protein backbone relaxation with 22 ns. The distribution of the different values also indicated some inherent side-chain flexibility within the Fingers and Palm domains, resulting in decreased values. Nevertheless, the quality of the measured data prevented detailed quantitative analysis at the current state providing only information for a sub-set of resonances.

The initial 2D [^13^C,^1^H]-NMR spectrum of MALVI^*proS*^-DinB had already shown some indications of specific line-broadening, which can possibly be attributed to conformational exchange processes. Therefore, we used a multiple quantum (MQ) Carr-Purcell-Meiboom-Gill (CPMG) relaxation dispersion experiment ^51^. We quantified the exchange-induced broadening effects (depicted as Δ*R*_2,eff_) on the micro- to millisecond timescale by measuring the difference in the relaxation rates at two different CPMG fields (25 and 750 Hz). The obtained values indicated that the vast majority of the methyl groups is not involved in chemical exchange processes as the values were close to zero in agreement with a stable protein fold (**Figure 3E, F**). Nevertheless, within the core of the Thumb domain several residues experience particular large exchange rates or were even exchange broadened, which is further proof of the existence of structural flexibility within this domain, that could possibly be attributed to movements of the different Thumb helices against each other within the core of this domain. In addition, some residues in the interface between Thumb and Palm also showed enhanced relaxation rates also pointing to a possible adaptation mechanism of the Thumb region in respect to the rest of the protein, which points to a possible allosteric signaling pathway originating from the Thumb domain towards the whole catalytic domain.

### DinB interaction with DNA

Hence, we next questioned on how the identified dynamics within the Thumb could contribute to DNA binding and how the PAD would achieve the postulated enclosing of the DNA. In order to investigate the DNA binding properties of DinB, we performed NMR titrations with a two complementary 18mer strands forming a double-stranded DNA (dsDNA), which was used previously in crystallographic studies of a DinB construct lacking the ten carboxy-terminal residues ^14^. Upon addition of the DNA, a distinct subset of the DinB backbone amide resonances exhibited signal intensity attenuations, clearly indicating the protein-DNA complex formation using a discrete set of DinB residues (**Figure 5A–C**). The observation of signal attenuation indicates, that on the chemical shift timescale, an interaction on the intermediate regime was observed with kinetic exchange rate constants ~100 – 1000 s^−1^, pointing to a dissociation constant in the low nanomolar range. Most of the residues undergoing intensity changes, are located at the supposed entry point of the DNA formed by the helices α8 and α11, which constitute parts of the Thumb domain oriented towards the inner cavity of the full-length DinB protein, as well as additional residues within the Palm and Fingers domains as well as the central β-sheet within the PAD comprising of strands β9–β12 (**Figure 5C**). We were especially intrigued by the observation of the effect on the Thumb helices, as these regions show enhanced pico– to nanosecond dynamics in the substrate-free state as described above. Thus, in the next step we employed the DinBΔPAD variant and repeated the titration experiment, showing a more localized and less pronounced effect mainly focused on the described two interaction points within the Thumb domain (**Figure 4D**). In contrast, using the isolated DinB-PAD did not show any significant chemical shift shift changes or signal intensity changes, therefore we concluded that this domain cannot bind the dsDNA in its isolated state. In summary a picture emerges, that the highly dynamic α8 and α11 helices of the Thumb domain are important for the binding of the DNA and might be even the determining interaction, and that the dsDNA once bound deeply into the groove encompassing Thumb, Palm, and Fingers regions the PAD finally encapsulates the DNA. Our data obtained here in solution are in excellent agreement with previously determined crystallographic studies ^14^, showing a deeply-bound DNA with the PAD domain enclosing it (**Figure 4E**).

**Figure 4.**
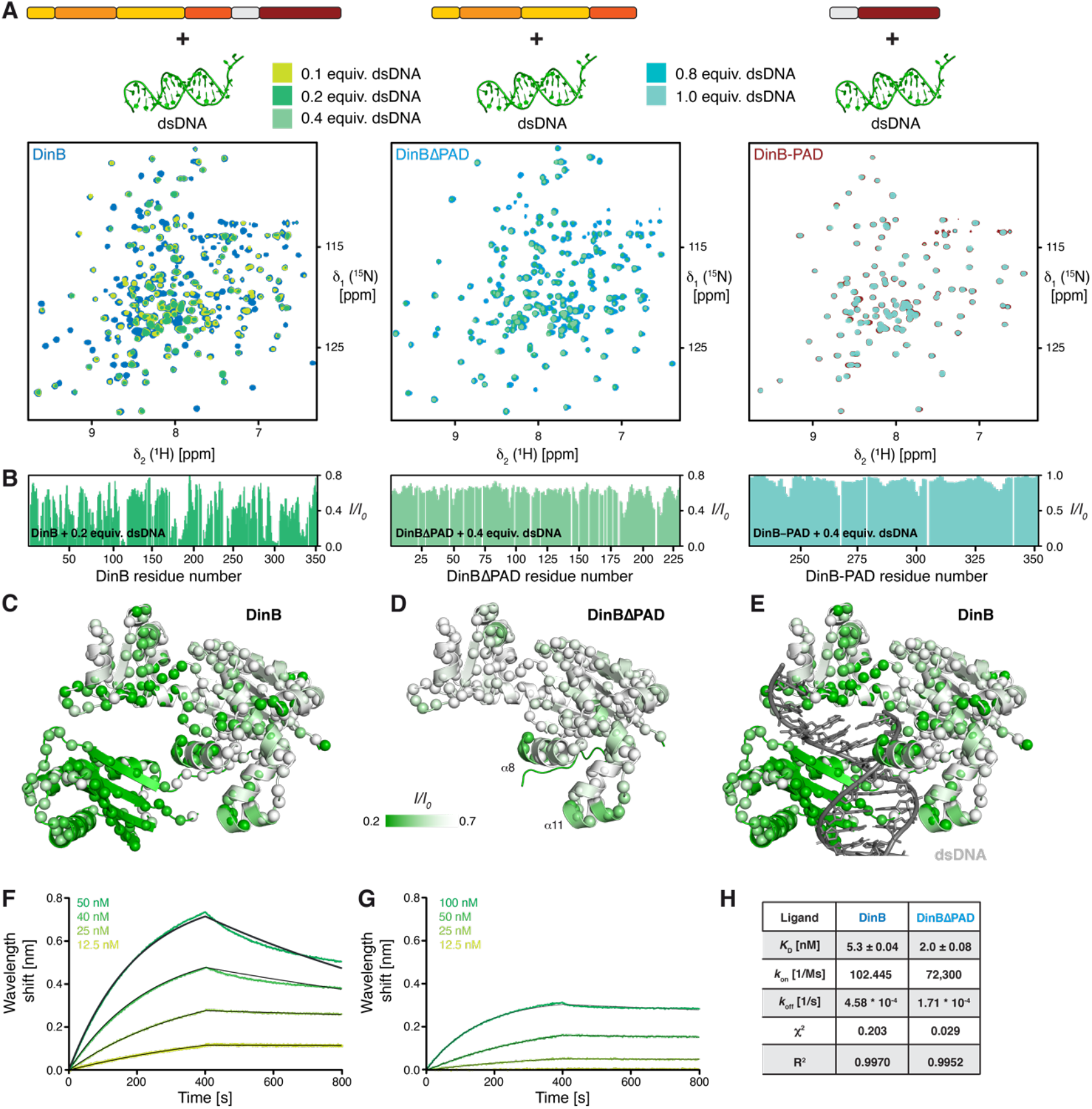
**A**) Titration of increasing amounts of dsDNA to either [*U*-^2^H,^15^N]-DinB, [*U*-^2^H,^15^N]-DinBΔPAD, or [*U*-^15^N]-DinB-PAD as indicated by the schematics on the top. Overlay of 2D [^15^N,^1^H]-NMR spectra of different DinB constructs in the absence (blue or red, respectively) and in the presence of increasing amounts of DNA as indicated by the green gradient acquired in NMR-buffer at 298 K. **B**) The ratio of the individual peak intensities in the presence of by the green gradient indicated equivalents of dsDNA to the respective apo DinB-forms plotted against the DinB residue number. **C, D**) Intensity changes upon dsDNA interaction were plotted on the DinB as well as the DinBΔPAD (PDB: 4Q45) structures by the indicated color gradient. The amide moieties of the individual construct are shown as spheres. **E**) Comparison of the obtained dsDNA interaction site for the full-length DinB obtained in solution with a previously determined DinB:dsDNA complex by X-ray crystallography (PDB: 4Q45) **F, G**) Bio-layer interferometry (BLI) data analysis of DinB (**G**) and DinBΔPAD (**F**) binding to dsDNA. Analyte concentrations are indicated. Non-linear least quare fits to the experimental data are indicated by the black lines. **H**) Table of the obtained dissociation constant *K*_D_, the kinetic parameters *k*_on_ and *k*_off_, as well as the χ^2^ and R^2^ parameters indicating the quality of the non-linear least squares fits.

**Figure 5.**
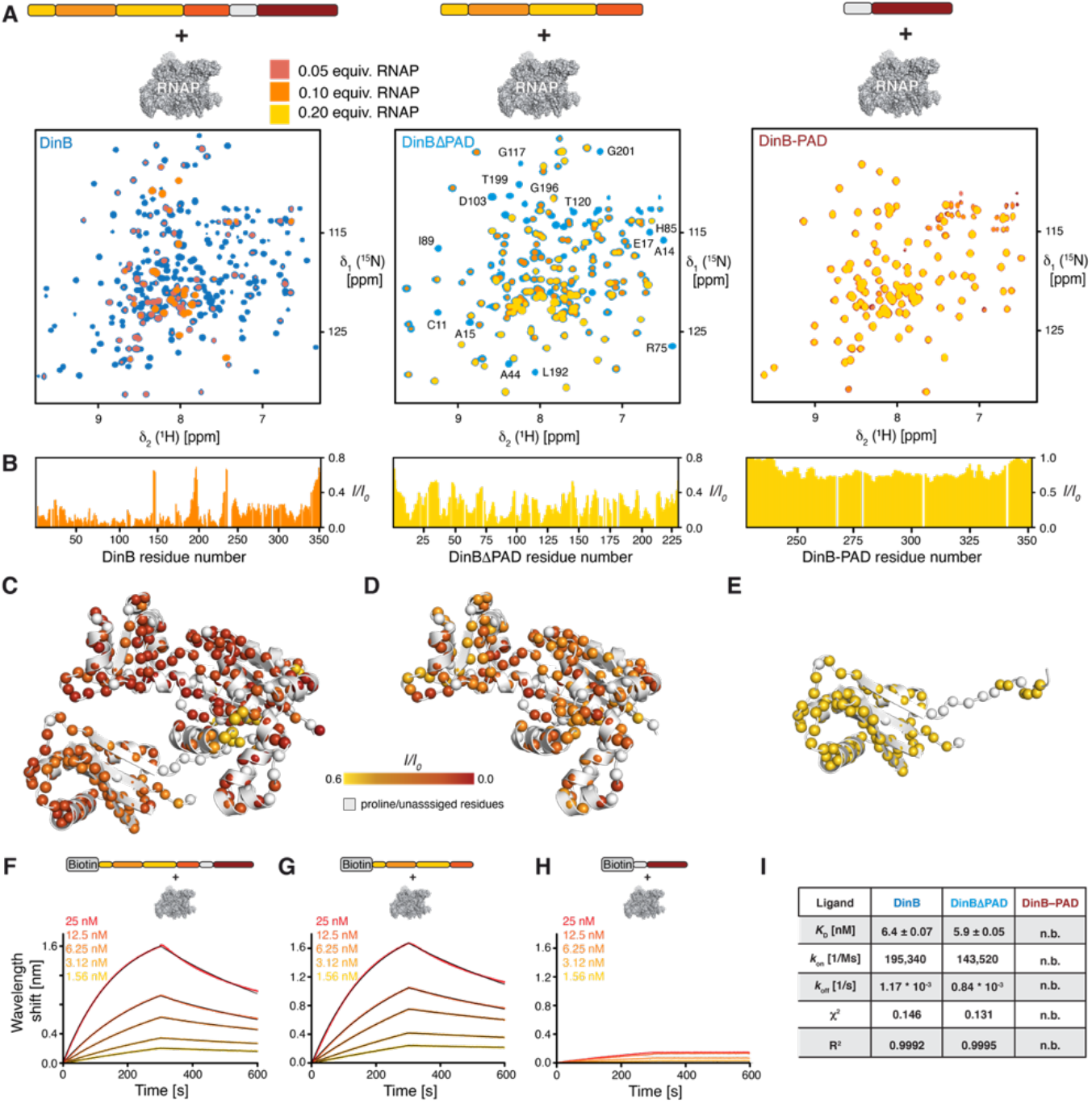
**A**) Titration of increasing amounts of RNAP to either [*U*-^2^H,^15^N]-DinB, [*U*-^2^H,^15^N]-DinBΔPAD, or [*U*-^15^N]-DinB-PAD as indicated by the schematics on the top. Overlay of 2D [^15^N,^1^H]-NMR spectra of different DinB constructs in the absence (blue or red, respectively) and in the presence of increasing amounts of RNAP as indicated by the orange to yellow gradient acquired in NMR-buffer at 298 K. **B**) The ratio of the individual peak intensities in the presence of by the orange to yellow gradient indicated equivalents of RNAP to the respective apo DinB-forms plotted against the DinB residue number. **C–E**) Intensity changes upon dsDNA interaction were plotted on the DinB as well as the DinBΔPAD and DinB-PAD (PDB: 4Q45) structures, respectively, by the indicated color gradient. The amide moieties of the individual construct are shown as spheres. **F–H**) BLI data analysis of DinB (**F**), DinBΔPAD (**G**), and DinB-PAD (**H**) binding to RNAP. Analyte concentrations are indicated. Non-linear least quare fits to the experimental data are indicated by the black lines. **I**) Table of the obtained dissociation constant *K*_D_, the kinetic parameters *k*_on_ and *k*_off_, as well as the χ^2^ and R^2^ parameters indicating the quality of the non-linear least squares fits. n.b. indicates non-binding.

Next, we used the BLI assay to quantitate the binding affinity of different DinB variants with the dsDNA. BLI data confirmed that DinB and DinBΔPAD bind the dsDNA with binding affinities for DinB (5.3 ± 0.04 nM) in comparison to DinBΔPAD (2.0 ± 0.08 nM) (**Figure 4F–H**), suggesting that the PAD domain does not contribute to the binding affinity in line with the observation of the absence of binding for the PAD in the BLI assay titrations, which is also in complete agreement with the observations in the NMR titration experiments. Furthermore, the observation of similar binding properties for the two larger constructs confirms our hypothesis that the initial binding event is the insertion of the helices α8 and α11 into the grooves of the dsDNA as also seen in the dsDNA:DinB complex structure (**Figure 4E**).

### DinB Palm and Finger regions facilitate RNAP binding

To address the RNAP-binding properties DinB, we initially performed NMR titrations adding unlabeled RNAP core-enzyme (comprising of the subunits α_2_ββ’ω) to [*U*-^2^H,^15^N]-DinB (**Figure 5A**). Already, upon addition of 0.1 molar ratio of RNAP, the backbone amide resonances of DinB exhibited severe line-broadening for a large number of resonances, owing the expected effect of the formation of a large DinB-RNAP complex with more than ~330 kDa in size, indicating a direct protein-protein interaction (**Figure 5B**). The effect was so strong that only a limited sub-set of resonances was clearly observable under these conditions mainly mapping to the unstructured parts of DinB as well as the PAD domain. To further elucidate the interaction in more detail, we employed the different constructs of DinB in a next step. Using the alternate DinBΔPAD construct clearly showed that this part of the protein is directly interacting with RNAP mainly through its Palm and Thumb region, in line with an extended binding interface. On the other hand, using the isolated DinB-PAD domain revealed only very minute signal intensity attenuations indicative for a transient and weak unspecific interaction in excellent agreement with the observation of the lowest degree of signal attenuation with this region within the full-length protein.

To assess the RNAP–DinB interaction in a more quantitative manner we used BLI analysis to characterize this interaction. The dissociation constant (*K*_D_) between DinB and the RNAP core-enzyme was determined to be 6.4 ± 0.07 nM (**Figure 5F**), whereas that of DinBΔPAD and the RNAP core-enzyme was found to be virtually identical with 5.9 ± 0.05 μM, with also only slight modulations of the on- and off-rates (**Figure 5G, I**). This finding together with the observation of no direct interaction with DinB-PAD (**Figure 5H**) indicative by the very limited wavelength shift of about a tenth compared to the other DinB constructs, as this change reports on the mass change on the BLI sensor even a larger change would be expected for such a small domain as observed in a previous study investigating RNAP interaction ^33^. Nevertheless, this observation is completely in line with the previous NMR titration experiments indicating only a transient and most likely unspecific interaction of RNAP with this domain.

## Discussion

Crystallographic studies of *E. coli* translesion DNA polymerase IV, DinB, revealed a right hand-folded catalytic core, which consists of three domains: Thumb, Fingers, and Palm and is connected to an additional polymerase-associated domain (PAD) *via* an extended flexible linker ^14^. DinB’s role in the adaptive stress-induced mutagenesis ^10^ is supposed to be governed by its inherent conformational changes in the catalytic core domains. Therefore, a detailed understanding of time-dependent motions of the domains is necessary to unravel the detailed molecular mechanism of its translesion DNA synthesis activity.

To this end we studied here DinB by solution NMR spectroscopy achieving almost complete resonance assignments of the protein backbone as well as the methyl bearing amino acid side chains. With this knowledge, we could for the first-time study in detail the substrate-free form of DinB, that eluded structural determination due to the flexibly attached PAD previous crystallization attempts as only the catalytic core of a DinB-homologue lacking the PAD could be crystallized ^58^. Overall, the structure in solution is almost identical to the crystal structures obtained in the complex with dsDNA (**Figure 1**) ^14^. Using a divide-and-conquer approach provided us with some additional insight about the Thumb and PAD domains. Whereas the isolated Thumb domain, consisting of five short α-helices, showed indications of a partially stable domain especially for the helices α7 and α8, clearly indicating that this domain needs further stabilization by additional interactions. In this case by an interaction surface on the PALM domain, which we could show leads to several inter-domain methyl-methyl NOEs (**Figure 2**) docking the Thumb onto the rest of the catalytic domain.

The PAD on the other hand forms a stable domain comprising of a four stranded β-sheet encompassed by two α-helices. Interestingly, despite being largely unfolded the linker between Thumb and PAD is ensuring the correct fold of the PAD as initial attempts to express and purify the isolated PAD domain lacking this linker resulted in unfolded protein (data not shown). This finding of the importance of an unstructured linker for the correct fold of the associated domain here matches previous finding for the CTD of bacterial UvrD, of helicase II ^33^. Based on computational studies in the case of Y-family DNA polymerases, it could be shown that the linker length and composition have a direct influence on the folding pathway of the whole DinB homologue studied ^59^, which agrees with our observation reported here. This finding bears of course the question, what is the functional role of this linker besides the folding process. Besides orienting the PAD domain at the right distance to bind a double stranded DNA as shown by previous crystallographic studies ^14,60,61^, another intriguing possibility is the effective concentration of this domain. It could be shown previously for unrelated enzymes, kinases that facilitate phosphorylation of their substrates, that in cases when the enzyme is tethered to the substrate these linker regions have a direct influence on the catalytic rate ^62,63^. In agreement with this general interpretation of the importance of the linker region already previous work on chimeric archaeal DinB homologous protein with varying linker sequences high-lighted the importance of this linker region for the catalytic efficiency of the enzyme ^15,26^. Furthermore, previous work could already show that the PAD domain is important for the enzymatic activity of DinB-type DNA polymerases stressing the important role of the correct positioning of this domain to ensure effective encapsulation of the DNA ^7^, which could also be interpreted in terms of the effective concentration needed for its inherent enzymatic activity ^62,64^.

Focusing subsequently on the backbone relaxation properties of full-length DinB showed in general a stable protein fold with a value 0.73 for the full-length protein, which in general indicates a stable fold, but is lower than the expected maximum values under the chosen experimental conditions of 0.86 ^65,66^ indicating the presence of global fast motions within the whole domain in solution. This analysis in particular pointed to significant flexibility of the Thumb domain as the average value for this region were even further decreased to 0.66, which together with the previously discussed indications of structural instability of this domain highlight the importance of structural and dynamical adaptations for this domain. In addition, our relaxation data also clearly shows that the uttermost carboxy-terminal residues show enhanced dynamics, which points to the lack of stable secondary structure elemnets in this region and thus in agreement with previous crystallographic studies employing a DinBΔC construct lacking the ten carboxy-terminal residues ^14^.

On the lower micro-second timescale, we observed a largely planar profile for the folded segment of the full-length protein, which is almost identical to that of the Thumb domain, and in agreement with the previous analysis of the fast timescale motions. Analyzing the fast and slow motions together revealed that the PAD is able to tumble to a certain extend independently of the catalytic core domain (**Supplementary Figure 1**), which is in agreement with previous study highlighting the functional importance of this feature ^59^.

Expanding the analysis of the inherent dynamics towards the methyl bearing amino-acid side chains we initially determined the product of the side-chain order parameters and the correlation time of the overall molecular tumbling (S^2^_axis_•τ_C_), which reports on the extend of the amplitude of motions on the fast NMR timescale gave a maximum value of ~23 ns, which is in good agreement with the τ_c_ for the protein backbone.

The obtained CPMG ΔR_2eff_ values indicate that the vast majority of the methyl groups is not involved in chemical exchange processes. Nevertheless, within the core of the Thumb domain in particular several residues experience large exchange rates or were even exchange broadened, which is further proof of the existence of structural flexibility within this domain and can possibly be attributed to conformational sampling to facilitate substrate interactions.

In line with this hypothesis, NMR titrations with dsDNA show a distinct subset of the DinB backbone amide resonances exhibiting signal intensity attenuations and thus binding to the DNA. Most of the residues undergoing intensity changes, are located at the supposed entry point of the DNA the helices α8 and α11 of the Thumb domain, parts of the Palm and Finger domains pointing to the inner cavity of the full-length DinB protein, and the central β-sheet within the PAD comprised of strands β9–β12. Thus, our data indicates that the initial recognition possibly occurs *via* helices α8 and α11 of the Thumb domain and then the dsDNA is bound on the inner cavity of the catalytic domain encompassing Palm and Fingers and in a last step locked in by the PAD-domain, that reaches around the dsDNA and enables optimal positioning ensuring efficient catalytic activity. The obtained binding affinities by BLI clearly confirm this as the DinBΔPAD construct shows almost the same affinity as the full-length DinB despite the NMR data showing a less extended binding interface (**Figure 4**).

In a last step we also examined if DinB can make direct contact with RNAP at a DNA lesion site, NMR titrations of unlabeled RNAP core-enzyme with [*U*-^2^H,^15^N]-DinB show the backbone amide resonances of DinB exhibited severe line-broadening for a large number of resonances, indicating a specific interaction and the formation of a large DinB-RNAP complex with more than ~330 kDa in size, and thus in line with previous observations for other enzymes involved in DNA repair ^33^. The obtained data points to an extended interaction surface on the outside of the DinB catalytic cleft mainly encompassing the Palm and Fingers domains, which was further evidenced by the almost abolished interaction for the isolated PAD domain (**Figure 5**). Functionally, this finding clearly points to the mechanistic possibility that DinB is still able to engage double stranded DNA while bound to the RNAP. The exact placing of the different proteins and eventual auxiliary binding partners, e.g. the transcription factor NusA, which has been implicated to bind to DinB ^67^, could be needed for the recruitment of DinB in the cellular context.

In summary, we show that the Thumb domain of DinB plays a central role in the binding of the DNA substrate and is modulated by inherent dynamics to exhibit conformational flexibility required for a stable initial interaction. The inherent dynamics are then transferred *via* allosteric signaling through the catalytic core as well as likely through the linker towards the PAD, thus ensuring a stable DNA encounter complex. Another crucial finding that warrants further investigation in the future is the direct interaction with the RNAP, which likely does not impair DNA engagement. Nevertheless, future structural studies are required to decipher the detailed mechanism of DinB function in the presence of the RNAP.

## Acknowledgements

The Swedish NMR Centre of the University of Gothenburg is acknowledged for spectrometer time. B.M.B. gratefully acknowledges funding from the Swedish Research Council (Starting Grant 2016-04721; Consolidator Grant 2020-00466) and the Knut och Alice Wallenberg Foundation through a Wallenberg Academy Fellowship (2016.0163 and 2020.0300) as well as through the Wallenberg Centre for Molecular and Translational Medicine, University of Gothenburg, Sweden. This study made use of NMRbox: National Center for Biomolecular NMR Data Processing and Analysis, a Biomedical Technology Research Resource (BTRR), which is supported by NIH grant P41GM111135 (NIGMS).

## Author Contributions

B.M.B. conceived the study and designed the experiments together with D.C.O.. D.C.O. with the support of I.M.-B. expressed and characterized protein variants. D.C.O. performed all other experimental work. D.C.O., J.L., and B.M.B. analyzed the data. All authors contributed ideas and discussions. D.C.O. and B.M.B. wrote jointly the manuscript with input from J.L..

## Competing Interests

The authors declare no conflict of interest.

## SUPPLEMENTARY INFORMATION

**Supplementary Table S1:**
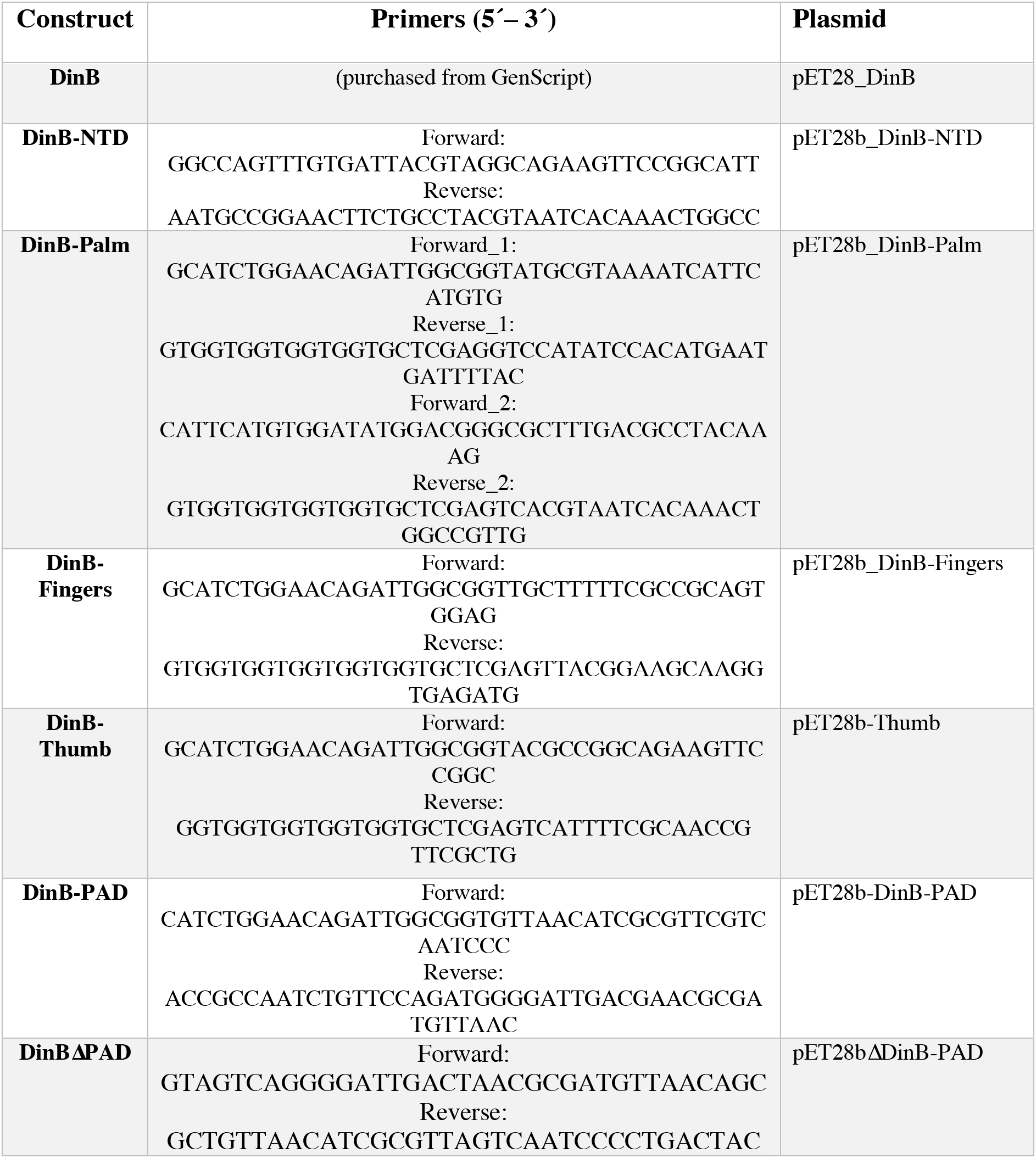
Primers used in this study to generate the different DinB constructs used.

**Supplementay Table S2:** Conditions used for the NMR experiments for the different DinB constructs used in this study.

| Protein | NMR buffer |
| --- | --- |
| **DinB** | 50 mM KPi, 300 mM KCl, 50 mM Arginine, 50 mM Glutamine, 1 mM EDTA, 1 mM TCEP, 1 mM CHAPSO, pH 6.5 |
| **DinBΔPAD** | PBS, 50 mM Arginine, 50 mM Glutamine, 1 mM EDTA, 1 mM TCEP, 0.1 mM CHAPSO, pH 7.4 |
| **DinB-PAD** | 50 mM KPi, 100 mM KCl, 50 mM Arginine, 50 mM Glutamine, 1 mM EDTA, 1 mM TCEP, 0.1 mM CHAPSO, pH 6.8 |
| **DinB-Thumb** | 50 mM KPi, 300 mM KCl, 50 mM Arginine, 50 mM Glutamine, 1 mM EDTA, 1 mM TCEP, 0.1 mM CHAPSO, pH 7.5 |

**Supplementary Figure S1.**
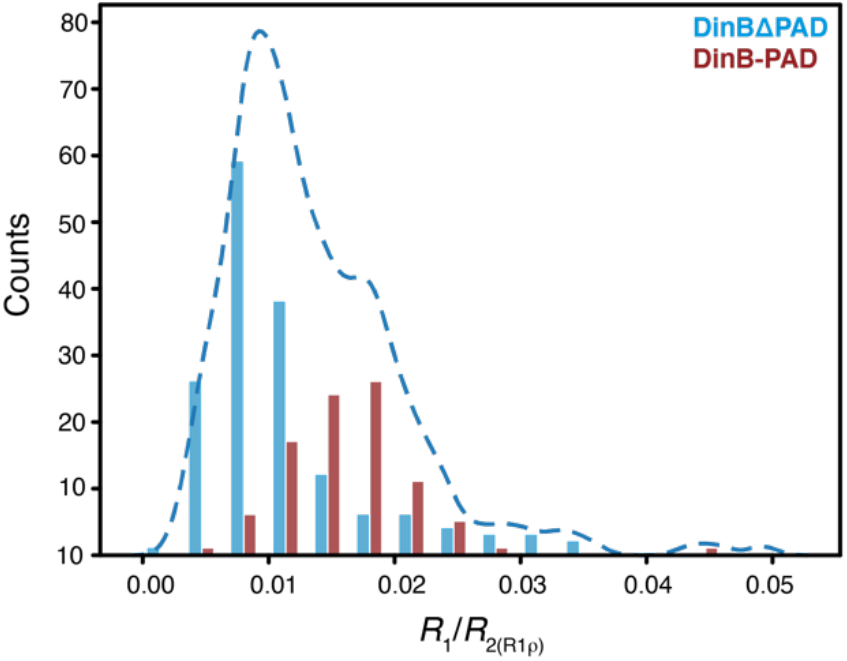
Histogram of *R*_1_/*R*_2(R1ρ)_ values of the DinBΔPAD (blue) and DinB-PAD (red). The dashed line corresponds to a kernel density estimation (KDE) of both domain values using a bandwidth of h=0.5. The bimodal distribution indicates partially decoupled motion of the DinB-PAD within full-length DinB.

**Supplementary Figure S2.**
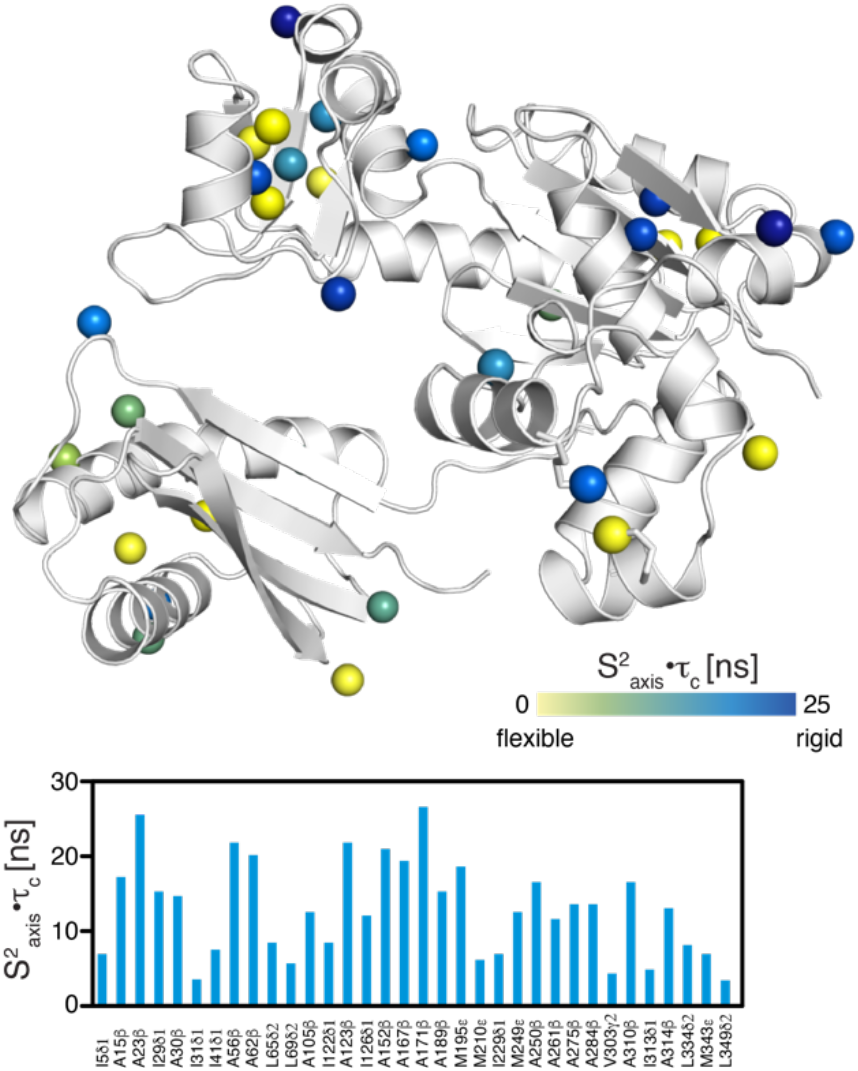
Local methyl group dynamics on the pico– to nanosecond timescale probed by methyl single-quantum (SQ) and triple-quantum (TQ) relaxation experiments showing the product of the local order parameter and the overall tumbling constant, S^2^_axis_ •τ_C_. the methyl groups are shown as spheres and the obtained S^2^_axis_•τ_C_-values by a yellow to blue gradient (top). S^2^_axis_•τ_C_-values plotted against the DinB amino acid sequence (bottom).

## Notes

### Competing Interest Statement

The authors have declared no competing interest.

### Summary of Updates

Figure 3 revised and updated.

## References

1. Klarer AC, McGregor WG. Replication of damaged genomes. Crit. Rev. in Eukaryot. Gene Expr. 21, 323–36 (2011).

2. Livneh Z, Shachar S. Multiple two-polymerase mechanisms in mammalian translesion DNA synthesis. Cell cycle. 9, 729–735 (2010).

3. Yang W, Woodgate R. What a difference a decade makes: Insights into translesion DNA synthesis. Proc. Natl. Acad. Sci. USA. 104, 15591–15598 (2007).

4. Friedberg EC, Fischhaber PL, Kisker C. Error-prone DNA polymerases: novel structures and the benefits of infidelity. Cell. 107, 9–12 (2001).

5. Ohmori H, Friedberg EC, Fuchs RP, et al. The Y-family of DNA polymerases. Mol. Cell. 8, 7–8 (2001).

6. Goodman MF. Error-prone repair DNA polymerases in prokaryotes and eukaryotes. Annu. Rev. Biochem. 71, 17–50 (2002).

7. Ling H, Boudsocq F, Woodgate R, Yang W. Crystal structure of a Y-family DNA polymerase in action: a mechanism for error-prone and lesion-bypass replication. Cell. 107, 91–102 (2001).

8. Silvian LF, Toth EA, Pham P, Goodman MF, Ellenberger T. Crystal structure of a DinB family error-prone DNA polymerase from *Sulfolobus solfataricus*. Nat. Struct. Biol.. 8, 984–989 (2001).

9. Trincao J, Johnson RE, Escalante CR, Prakash S, Prakash L, Aggarwal AK. Structure of the catalytic core of *S. cerevisiae* DNA polymerase η: implications for translesion DNA synthesis. Mol. Cell. 8, 417–426 (2001).

10. McKenzie GJ, Lee PL, Lombardo MJ, Hastings P, Rosenberg SM. SOS mutator DNA polymerase IV functions in adaptive mutation and not adaptive amplification. Mol. Cell. 7, 571–579 (2001).

11. Kenyon CJ, Walker GC. DNA-damaging agents stimulate gene expression at specific loci in *Escherichia coli*. Proc. Natl. Acad. Sci. USA. 77, 2819–2823 (1980).

12. Shen X, Sayer JM, Kroth H, et al. Efficiency and accuracy of SOS-induced DNA polymerases replicating benzo [α] pyrene-7,8-diol 9,10-epoxide A and G adducts. J. Biol. Chem. 277, 5265–5274 (2002).

13. Tang M, Pham P, Shen X, et al. Roles of *E. coli* DNA polymerases IV and V in lesion-targeted and untargeted SOS mutagenesis. Nature. 404, 1014–1018 (2000).

14. Kottur J, Sharma A, Gore KR, et al. Unique structural features in DNA polymerase IV enable efficient bypass of the N^2^ adduct induced by the nitrofurazone antibiotic. Structure. 23, 56–67 (2015).

15. Boudsocq F, Kokoska RJ, Plosky BS, et al. Investigating the role of the little finger domain of Y-family DNA polymerases in low fidelity synthesis and translesion replication. J. Biol. Chem. 279, 32932–32940 (2004).

16. Yang W. An overview of Y-family DNA polymerases and a case study of human DNA polymerase η. Biochemistry. 53, 2793–2803 (2014).

17. Bunting KA, Roe SM, Pearl LH. Structural basis for recruitment of translesion DNA polymerase Pol IV/DinB to the β-clamp. EMBO J. 22, 5883–5892 (2003).

18. Cohen SE, Walker GC. New discoveries linking transcription to DNA repair and damage tolerance pathways. Transcription 2, 37–40 (2011).

19. Heltzel JMH, Maul RW, Scouten Ponticelli SK, Sutton MD. A model for DNA polymerase switching involving a single cleft and the rim of the sliding clamp. Proc. Natl. Acad. Sci. USA. 106, 12664–12669 (2009).

20. Heltzel JMH, Maul RW, Wolff DW, Sutton MD. *Escherichia coli* DNA polymerase IV (Pol IV), but not Pol II, dynamically switches with a stalled Pol III* replicase. J. Bacteriol. 194, 3589–3600 (2012).

21. Cohen SE, Godoy VG, Walker GC. Transcriptional modulator NusA interacts with translesion DNA polymerases in *Escherichia coli*. J. Bacteriol. 191, 665–672 (2009).

22. Brotcorne-Lannoye A, Maenhaut-Michel G. Role of RecA protein in untargeted UV mutagenesis of bacteriophage λ: evidence for the requirement for the *dinB* gene. Proc. Natl. Acad. Sci. USA. 83, 3904–3908 (1986).

23. Donnelly CE, Walker GC. Coexpression of UmuD’with UmuC suppresses the UV mutagenesis deficiency of *groE* mutants. J. Bacteriol. 174, 3133–3139 (1992).

24. Wagner J, Fujii S, Gruz P, Nohmi T, Fuchs RP. The β clamp targets DNA polymerase IV to DNA and strongly increases its processivity. EMBO Rep. 1, 484–488 (2000).

25. Uljon SN, Johnson RE, Edwards TA, Prakash S, Prakash L, Aggarwal AK. Crystal structure of the catalytic core of human DNA polymerase κ. Structure. 12, 1395–1404 (2004).

26. Wilson RC, Jackson MA, Pata JD. Y-family polymerase conformation is a major determinant of fidelity and translesion specificity. Structure. 21, 20–31 (2013).

27. Mikolajczyk J, Drag M, Békés M, Cao JT, Ronai Z, Salvesen GS. Small ubiquitin-related modifier (SUMO)-specific proteases: profiling the specificities and activities of human SENPs. J. Biol. Chem. 282, 26217–26224 (2007).

28. Azatian SB, Kaur N, Latham MP. Increasing the buffering capacity of minimal media leads to higher protein yield. J. Biomol. NMR. 73,11–17 (2019).

29. Gans P, Hamelin O, Sounier R, et al. Stereospecific isotopic labeling of methyl groups for NMR spectroscopic studies of high-molecular-weight proteins. Angew. Chem. Int. Ed. Engl. 49, 1958–1962 (2010).

30. Callon M, Burmann BM, Hiller S. Structural mapping of a chaperone–substrate interaction surface. Angew. Chem. Int. Ed. Engl. 53, 5069–5072 (2014).

31. Goto NK, Gardner KH, Mueller GA, Willis RC, Kay LE. A robust and cost-effective method for the production of Val, Leu, Ile (δ_1_) methyl-protonated ^15^N-, ^13^C-, ^2^H-labeled proteins. J. Biomol. NMR. 13, 369–374 (1999).

32. Svetlov V, Artsimovitch I. Purification of bacterial RNA polymerase: tools and protocols. Methods Mol. Biol. 1276, 13–29 (2015).

33. Kawale AA, Burmann BM. UvrD helicase–RNA polymerase interactions are governed by UvrD’s carboxy-terminal Tudor domain. Commun. Biol. 3, 607 (2020).

34. Pervushin K, Riek R, Wider G, Wüthrich K. Attenuated T_2_ relaxation by mutual cancellation of dipole–dipole coupling and chemical shift anisotropy indicates an avenue to NMR structures of very large biological macromolecules in solution. Proc. Natl. Acad. Sci. USA. 94, 12366–12371 (1997).

35. Salzmann M, Pervushin K, Wider G, Senn H, Wüthrich K. TROSY in triple-resonance experiments: New perspectives for sequential NMR assignment of large proteins. Proc Natl Acad Sci USA. 95, 13585–13590 (1998).

36. Sattler M, Schleucher J, Griesinger C. Heteronuclear multidimensional NMR experiments for the structure determination of proteins in solution. Prog. Nucl. Magn. Reson. Spectrosc. 34, 93–158 (1999).

37. Rossi P, Xia Y, Khanra N, Veglia G, Kalodimos CG. ^15^N and ^13^C-SOFAST-HMQC editing enhances 3D-NOESY sensitivity in highly deuterated, selectively [^1^H,^13^C]-labeled proteins. J. Biomol. NMR. 66, 259–271 (2016).

38. Wider G, Dreier L. Measuring Protein concentrations by NMR spectroscopy. J. Am. Chem. Soc. 128, 2571–2576 (2006).

39. Delaglio F, Grzesiek S, Vuister GW, Zhu G, Pfeifer J, Bax A. NMRPipe: a multidimensional spectral processing system based on UNIX pipes. J. Biomol. NMR. 6, 277–293 (1995).

40. Jaravine V, Ibraghimov I, Orekhov VY. Removal of a time barrier for high-resolution multidimensional NMR spectroscopy. Nat. Methods. 3, 605–607 (2006).

41. Keller R. The Computer-Aided Resonance Assignment Tutorial (Cantina Verlag, Goldau, 2004).

42. Nielsen JT, Mulder FAA. POTENCI: prediction of temperature, neighbor and pH-corrected chemical shifts for intrinsically disordered proteins. J. Biomol. NMR. 70, 141–165 (2018).

43. Morgado L, Burmann BM, Sharpe T, Mazur A, Hiller S. The dynamic dimer structure of the chaperone Trigger Factor. Nat. Commun. 8, 1992 (2017).

44. Burmann BM, Wang C, Hiller S. Conformation and dynamics of the periplasmic membrane-protein–chaperone complexes OmpX–Skp and tOmpA–Skp. Nat. Struc. Mol. Biol. 20, 1265–1272 (2013).

45. Lakomek NA, Ying J, Bax A. Measurement of ^15^N relaxation rates in perdeuterated proteins by TROSY-based methods. J. Biomol. NMR. 53, 209–221 (2012).

46. Ahlner A, Carlsson M, Jonsson BH, Lundström P. PINT: a software for integration of peak volumes and extraction of relaxation rates. J. Biomol. NMR. 56, 191–202 (2013).

47. Dosset P, Hus JC, Marion D, Blackledge M. A novel interactive tool for rigid-body modeling of multi-domain macromolecules using residual dipolar couplings. J. Biomol. NMR. 20, 223–231 (2001).

48. Maciejewski MW, Schuyler AD, Gryk MR, et al. NMRbox: A Resource for biomolecular NMR computation. Biophys. J. 112, 1529–1534 (2017).

49. Sun H, Kay LE, Tugarinov V. An optimized relaxation-based coherence transfer NMR experiment for the measurement of side-chain order in methyl-protonated, highly deuterated proteins. J. Phys. Chem. B. 115, 14878–14884 (2011).

50. Weinhäupl K, Lindau C, Hessel A, et al. Structural basis of membrane protein chaperoning through the mitochondrial ïntermembrane space. Cell. 175, 1365–1379 (2018).

51. Korzhnev DM, Kloiber K, Kanelis V, Tugarinov V, Kay LE. Probing slow dynamics in high molecular weight proteins by methyl-TROSY NMR spectroscopy: Application to a 723-residue enzyme. J Am Chem Soc. 2004;126(12):3964–3973. doi:10.1021/ja039587i

52. Burmann BM, Gerez JA, Matečko-Burmann I, et al. Regulation of α-synuclein by chaperones in mammalian cells. Nature. 577, 127–132 (2020).

53. Selo I, Négroni L, Créminon C, Grassi J, Wal J. Preferential labeling of α-amino N-terminal groups in peptides by biotin: application to the detection of specific anti-peptide antibodies by enzyme immunoassays. J. Immunol. Methods. 199, 127–138 (1996).

54. Moro SL, Cocco MJ. ^1^H, ^13^C, and ^15^N backbone resonance assignments of the full-length 40 kDa *S. acidocaldarius* Y-family DNA polymerase, dinB homolog. Biomol. NMR Assign. 9, 441–445 (2015).

55. Palmer 3rd A, Kroenke CD, Loria JP. Nuclear magnetic resonance methods for quantifying microsecond-to-millisecond motions in biological macromolecules. Meth. Enzymol. 339, 204–238 (2001).

56. Burmann BM, Knauer SH, Sevostyanova A, et al. An α helix to β barrel domain switch transforms the transcription factor RfaH into a translation factor. Cell. 150, 291–303 (2012).

57. Burmann BM, Scheckenhofer U, Schweimer K, Rösch P. Domain interactions of the transcription–translation coupling factor *Escherichia coli* NusG are intermolecular and transient. Biochem. J. 435, 783–789 (2011).

58. Zhou BL, Pata JD, Steitz TA. Crystal structure of a DinB lesion bypass DNA polymerase catalytic fragment reveals a classic polymerase catalytic domain. Mol. Cell. 8, 427–437 (2001).

59. Chu X, Suo Z, Wang J. Investigating the trade-off between folding and function in a multidomain Y-family DNA polymerase. Elife. 9, e60434 (2020).

60. Kottur J, Nair DT. Pyrophosphate hydrolysis is an intrinsic and critical step of the DNA synthesis reaction. Nucleic Acids Res. 46, 5875–5885 (2018).

61. Sharma A, Kottur J, Narayanan N, Nair DT. A strategically located serine residue is critical for the mutator activity of DNA polymerase IV from *Escherichia coli*. Nucleic acids research. 41, 5104–5114 (2013).

62. Dyla M, Kjaergaard M. Intrinsically disordered linkers control tethered kinases *via* effective concentration. Proc. Natl. Acad. Sci. USA. 117, 21413–21419 (2020).

63. Kjaergaard M. Estimation of effective concentrations enforced by complex linker architectures from conformational ensembles. Biochemistry. 61, 171–182 (2022).

64. Dyla M, González Foutel NS, Otzen DE, Kjaergaard M. The optimal docking strength for reversibly tethered kinases. Proc. Natl. Acad. Sci. USA. 119, e2203098119 (2022).

65. Kay LE, Torchia DA, Bax A. Backbone dynamics of proteins as studied by nitrogen-15 inverse detected heteronuclear NMR spectroscopy: application to staphylococcal nuclease. Biochemistry. 28, 8972–8979 (1989).

66. Kay LE. NMR studies of protein structure and dynamics. J. Magn. Reson. 173, 193–207 (2005).

67. Cohen SE, Lewis CA, Mooney RA, et al. Roles for the transcription elongation factor NusA in both DNA repair and damage tolerance pathways in *Escherichia coli*. Proc. Natl. Acad. Sci. USA. 107, 15517–15522 (2010).

